# APOBEC3A/B-induced mutagenesis is responsible for 20% of heritable mutations in the TpCpW context

**DOI:** 10.1101/054197

**Authors:** Vladimir B. Seplyarskiy, Maria A. Andrianova, Georgii A. Bazykin

## Abstract

APOBEC3A/B cytidine deaminase is responsible for the majority of cancerous mutations in a large fraction of cancer samples. However, its role in heritable mutagenesis remains very poorly understood. Recent studies have demonstrated that both in yeast and in human cancerous cells, most of APOBEC3A/B-induced mutations occur on the lagging strand during replication. Here, we use data on rare human polymorphisms, interspecies divergence, and de novo mutations to study germline mutagenesis, and analyze mutations at nucleotide contexts prone to attack by APOBEC3A/B. We show that such mutations occur preferentially on the lagging strand. Moreover, we demonstrate that APOBEC3A/B-like mutations tend to produce strand-coordinated clusters, which are also biased towards the lagging strand. Finally, we show that the mutation rate is increased 3’ of C→G mutations to a greater extent than 3’ of C→T mutations, suggesting pervasive translesion bypass of the APOBEC3A/B-induced damage. Our study demonstrates that 20% of C→T and C→G mutations segregating as polymorphisms in human population are attributable to APOBEC3A/B activity.

## Introduction

Understanding the processes responsible for heritable mutations is important for a broad range of evolutionary, population genetics and medical questions^1^. Recent studies have shown that different numbers of *de novo* mutations are inherited from father and from mother, with the contribution of paternal mutations being 2-4 times higher^2^^−^^4^. Additionally, the number of *de novo* mutations strongly depends on father’s age^2^^−^^4^ and, to a lesser extent, on mother’s age at conception^4^. At the molecular level, the understanding of mechanisms of heritable mutagenesis is very limited. Only a few of the mutation types can be attributed to specific molecular processes. Most prominently, the <u>C</u>pG→TpG substitutions are known to result from spontaneous cytosine deamination of methyl-cytosine in the <u>C</u>pG context together with poor efficiency of subsequent base excision repair^5^,^6^; mutations in the CpCp<u>C</u> motif arise due to activity of the APOBEC3G protein which is normally involved in protection against viruses and retroelements^7^,^8^; and small insertions and deletions result from polymerase slippage on homonucleotide tracts and tandem repeats^9^.

The sources of somatic mutations, in particular those in cancers, are better understood. The rates of such mutations were related to age-dependent cytosine deamination, to activity of APOBEC3A/B/G and AID, to deficiencies in systems responsible for the fidelity of DNA repair and replication and to exposure to external and internal mutagens^10^,^11^.

APOBEC3A/B-induced mutations were described for many cancer types^10^,^12^,^13^. Mutations produced by APOBEC3A/B have known properties confirmed both in yeast and in human cancers: (i) they are C→D mutations in the Tp<u>C</u>pN context (the more specific APOBEC3A/B signature is Tp<u>C</u>pW→K, where D denotes A, T or G, W denotes A or T and K denotes G or T)^14^–^17^; (ii) they often form strand-coordinated clusters^12^,^16^,^18^; and (iii) they are strongly biased towards the lagging strand during replication^14^,^19^,^20^. Finally (iv), cytosines deaminated to uracils by APOBEC frequently result in C→G substitutions. According to the current models, these mutations arise due to incomplete repair of U-G mismatches resulting in abasic sites. In turn, abasic sites are bypassed by REV1, which inserts exactly one nucleotide opposite to the site, and this single nucleotide primer is then extended by the low fidelity polymerase ζ ^18^,^21^,^22^. Additionally, as we show here, (v) strand coordinated clusters in cancers are very strongly biased towards the lagging strand.

Here, by determining the prevalent direction of the replication fork at each genomic region, we show that APOBEC3A/B contributes substantially to heritable mutagenesis. We show that Tp<u>C</u>pW→K mutations in human polymorphism and divergence exhibit all of the above properties of APOBEC3A/B-induced mutations, unlike non-APOBEC-induced Vp<u>C</u>pW→K mutations (where V denotes A, C or G). We estimate that about 20% of heritable Tp<u>C</u>pW→K mutations, or 1.4% of all heritable mutations, are linked to APOBEC3A/B activity.

## Results

### APOBEC3A/B-induced mutations in cancer samples are 3 times more frequent on the lagging strand

We previously demonstrated that fork polarity (fp) calculated as the derivative of replication timing allows to discriminate between the strands replicated as leading vs. as lagging^14^. In tissue-matched data on human cancers, the rate of Tp<u>C</u>pW→K mutations on the lagging strand is ~3 times higher than that on the leading strand (Figure S1); this bias is stronger than that previously observed using replication timing data from a different tissue^14^,^19^. Therefore, accumulation of APOBEC3A/B mutations in human cancers is strongly strand-specific. We asked if this specificity, as well as other properties of APOBEC3A/B-induced mutagenesis, are also manifested in heritable mutations.

### Heritable mutations in the APOBEC3A/B context are 20% more frequent on the lagging strand

Rare polymorphisms tend to be young. Therefore, their mutational spectra and genomic distribution much better reflect the spectra of *de novo* mutations, compared to the spectra and distribution of substitutions between species which are affected by non-mutational processes such as selection or biased gene conversion operating over the lifetime of a mutation as it spreads to fixation^23^,^24^. Therefore, to study heritable mutations, we mainly focused on rare polymorphisms^25^, i.e. those with the frequency of the derived allele in the human population below 1%.

We measured the ratio of the frequencies of C→K mutations to those of complementary G→M (where M denotes A or C) mutations in different contexts, and asked how this ratio depends on whether the analyzed strand is preferentially replicated as leading or as lagging. The rates of both provisionally APOBEC3A/B-induced mutation types (to T and to G) in both APOBEC3A/B contexts (Tp<u>C</u>pT and Tp<u>C</u>pA) were 10-20% higher on the lagging strand (Figure 1), while a weak bias was observed in only one of the comparisons for the corresponding mutations in the non-APOBEC3A/B Vp<u>C</u>pW context (Figure 1A). This implies that ~15-30% of Tp<u>C</u>pW→K substitutions resulted from APOBEC3A/B deamination (see Methods for details), with a higher fraction for the mutations in the Tp<u>C</u>pT context (~30%) than in the Tp<u>C</u>pA context (~15%). Similar, albeit slightly weaker, biases were observed for interspecies differences accumulated in the human lineage since the last common ancestor with chimpanzee (Figure S2).

**Figure 1.**
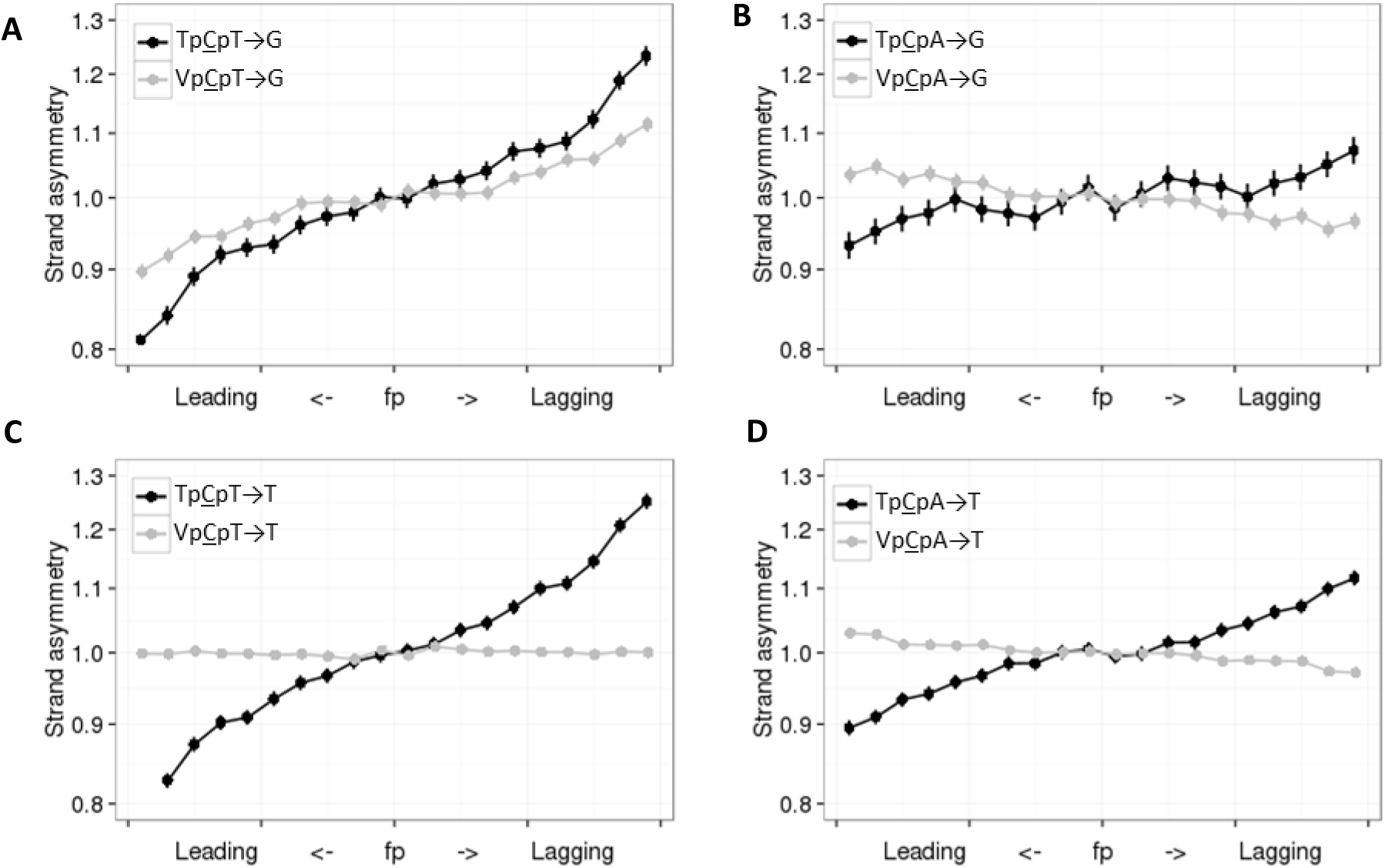
Mutations in APOBEC3A/B context are more frequent on the lagging strand. Horizontal axis, propensity of the region of the DNA strand to be replicated as lagging or leading. Vertical axis, ratio of the frequencies of the two complementary mutation types on the strand in this category. Vertical bars represent 95% confidence intervals. V corresponds to A, C or G.

APOBEC3A and APOBEC3B-induced mutations preferentially occur when respectively Y (T or C) or R (A or G) is observed two nucleotides upstream of the mutated C^16^,^17^. Therefore, by analyzing this extended context, we are able to discriminate between these two enzymes. The strand bias is stronger for YpTp<u>C</u>pW contexts than for RpTp<u>C</u>pW contexts (Figure S3), suggesting that, similarly to cancers with a high burden of APOBEC-induced mutations^17^, APOBEC3A likely contributes more to the observed mutations than APOBEC3B.

In cancers most affected by APOBEC3A/B-induced mutagenesis, the mutation rate in the Tp<u>C</u>pW context is increased by up to 40 fold^14^. From strand asymmetry, we estimate that only 15-30% of heritable Tp<u>C</u>pW→K mutations are induced by APOBEC3A/B. We asked whether this APOBEC3A/B-induced mutagenesis is also manifested in an increased genome-wide mutation rate in the corresponding nucleotide context. However, the frequencies of Tp<u>C</u>pW→K were not uniformly higher than the frequencies of Vp<u>C</u>pW→K mutations (Table S1). This is probably due to a stronger relative contribution on APOBEC3A/B-independent context preferences^26^ to polymorphism data, compared to cancer samples where APOBEC3A/B virtually monopolizes the mutation process.

### Heritable mutations in APOBEC3A/B context tend to form strand coordinated clusters

As APOBEC family enzymes deaminate single stranded DNA, they tend to produce strand coordinated clusters. To study such clusters, we focused on pairs of rare single nucleotide polymorphisms (SNPs) in strong linkage at distances of up to 5000 nucleotides from each other (see Methods for details). Mutations at sites closer than 10 nucleotides to each other occur due to activity of low fidelity polymerases or other mechanisms not related to APOBEC3A/B^27^^−^^29^; therefore, we excluded such pairs from our analysis. Over a half of remaining pairs of Tp<u>C</u>pW→K mutations were strand coordinated, and for three out of the four considered mutation types, the fraction of strand coordinated pairs was significantly higher than for the same mutations in the Vp<u>C</u>pW context used as a control (Figure 2A-D). Similar results were obtained for interspecies divergence data (Table S2).

**Figure 2.**
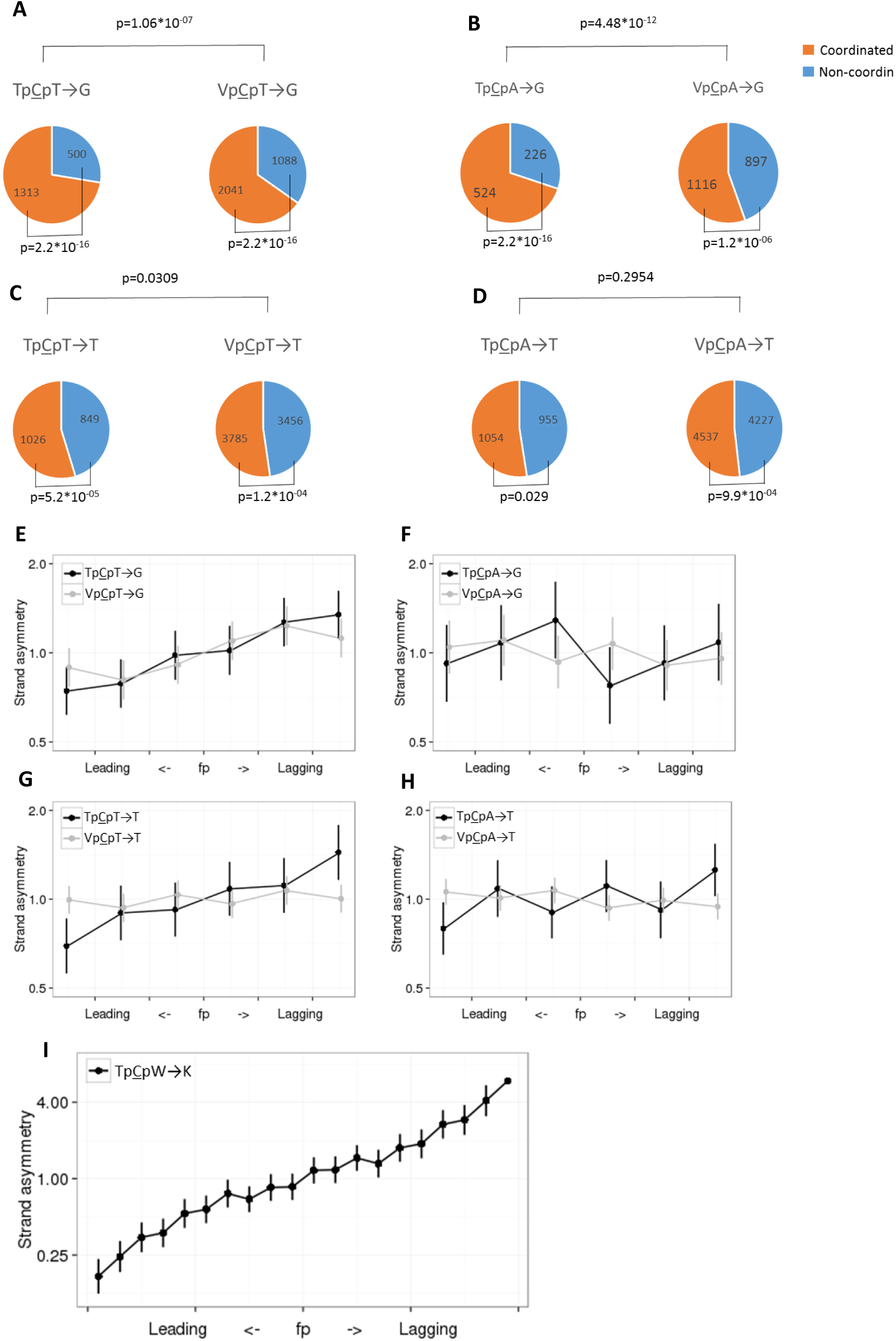
Clustered mutations in the Tp<u>C</u>pW→K context are enriched in strand coordinated clusters and biased towards the lagging strand. **(A-D)** Pairs of linked SNPs in contexts prone to APOBEC3A/B-induced mutations tend to be strand coordinated. **(E-H)** Such strand coordinated pairs are biased towards the lagging strand. **(I)** Strand bias in strand coordinated clusters in breast cancer samples with the APOBEC3A/B mutational signature. In (E-I), the axes and notation are as in Figure 1.

APOBEC3A/B-induced mutations are biased towards the lagging strand in cancer and in yeasts. We asked whether this pattern is also observed for strand-coordinated clusters. In cancers with a strong prevalence of APOBEC3A/B-related mutagenesis, strand-coordinated clusters occur six fold more frequently on the lagging strand, demonstrating the highest level of replicative strand asymmetry reported for mammalian cells (Figure 2I). Therefore, we could expect an excess of strand-coordinated pairs of heritable Tp<u>C</u>pW→K mutations on the lagging strand if APOBEC3A/B plays a role in their formation. Indeed, clustered mutations preferentially occur on the lagging strand (Figure 2E-H), and the level of strand asymmetry for them is higher than that for dispersed mutations (Figure S4), reflecting an enrichment of APOBEC3A/B-induced clusters among strand coordinated clusters.

### An unknown mechanism produces clusters of heritable Np<u>C</u>pT→G mutations

Notably, a tendency to form strand-coordinated clusters was observed not only for the Tp<u>C</u>pW→K mutations, but, to a lesser extent, also for the Vp<u>C</u>pW→K mutations (Figure 2A-D). To better understand this, we focused on the Vp<u>C</u>pT→G mutations for which this trend is particularly strong (Figure 2C), and which are also biased towards the lagging strand (Figure 1A). Notably, C→G is among the most frequent mutations in *de novo* mutational clusters^3^. We investigated C→G mutations in more detail in two recently published whole-genome datasets on human trios^3^,^4^ and found that within 23 pairs of C→G mutations at distances of up to 20Kb from each other, 18 were strand coordinated, which is more than expected if they were independent (p=0.01; Figure 3A). The context preferences of the clustered C→G mutations differed from those of non-clustered mutations (p=0.019), with the clustered C→G mutations enriched in the Np<u>C</u>pT→G context (p=0.04, Figure 3C). Therefore, while the mechanism behind the strand coordinated Np<u>C</u>pT→G mutations remains unclear, they are observed in all data types.

**Figure 3.**
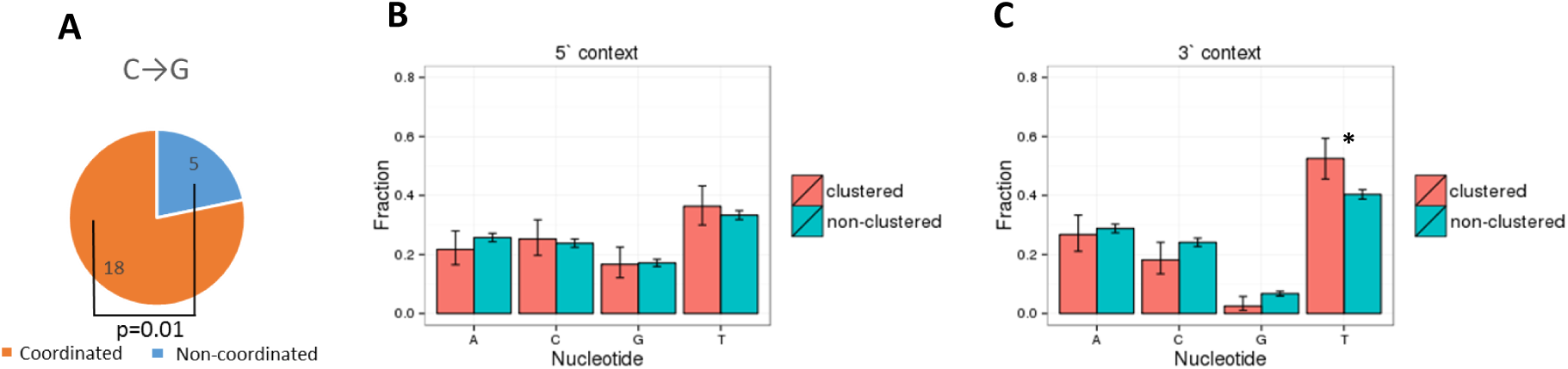
Properties of C→G mutations involved in clusters of *de novo* mutations. **(A)** Pairs of C→G *de novo* mutations tend to form strand coordinated clusters. **(B, C)** Fractions of different nucleotides at adjacent sites 5` **(B)** and 3` **(C)** of the mutated position calculated for clustered and dispersed C→G *de novo* mutations. Vertical bars represent 95% confidence intervals. * p < 0.05 (chi-square test).

### REV1 and polymerase ζ cause C→G SNPs and increase the mutation rate 3’ of them

In cancers, up to a half of APOBEC3A/B-induced mutations are C→G. These mutations arise due to bypass of abasic sites that originate from unfinished repair of cytosines deaminated by APOBEC3A/B. Abasic sites cause stalling of replicative polymerases and replication fork arrest. Two enzymes involved in bypass of abasic sites are REV1 which introduces a single-nucleotide primer (causing the C→G mutation itself) and low fidelity polymerase ζ which replicates a stretch of DNA downstream of it^18^. If similar processes affect heritable mutations, we expect to observe a higher mutation rate downstream (3’) of the C→G mutations that mark the start of polymerase ζ-dependent DNA synthesis. We compared the mutation rates at distances of up to 5Kb 5’ and 3’ of Tp<u>C</u>pW→K mutations. The mutation rates were increased ~1-2Kb 3’ ofC→G mutations in the APOBEC3A/B contexts both on the leading and the lagging strand (Figure 4A, C, E, G). A higher mutation rate 3’ of Tp<u>C</u>pW→G SNPs on both replicative strands at distances of up to 2kb is in agreement with the key role of polymerase ζ in generation of these mutations, and with its known low (~1Kb) processivity^30^. However, an increase in the mutation rate is also observed 3’ of Vp<u>C</u>pW→G mutations (Figure S5), implying that the role of polymerase ζ in the accumulation of heritable C→G mutations is not limited to bypass of APOBEC3A/B-induced abasic sites.

**Figure 4.**
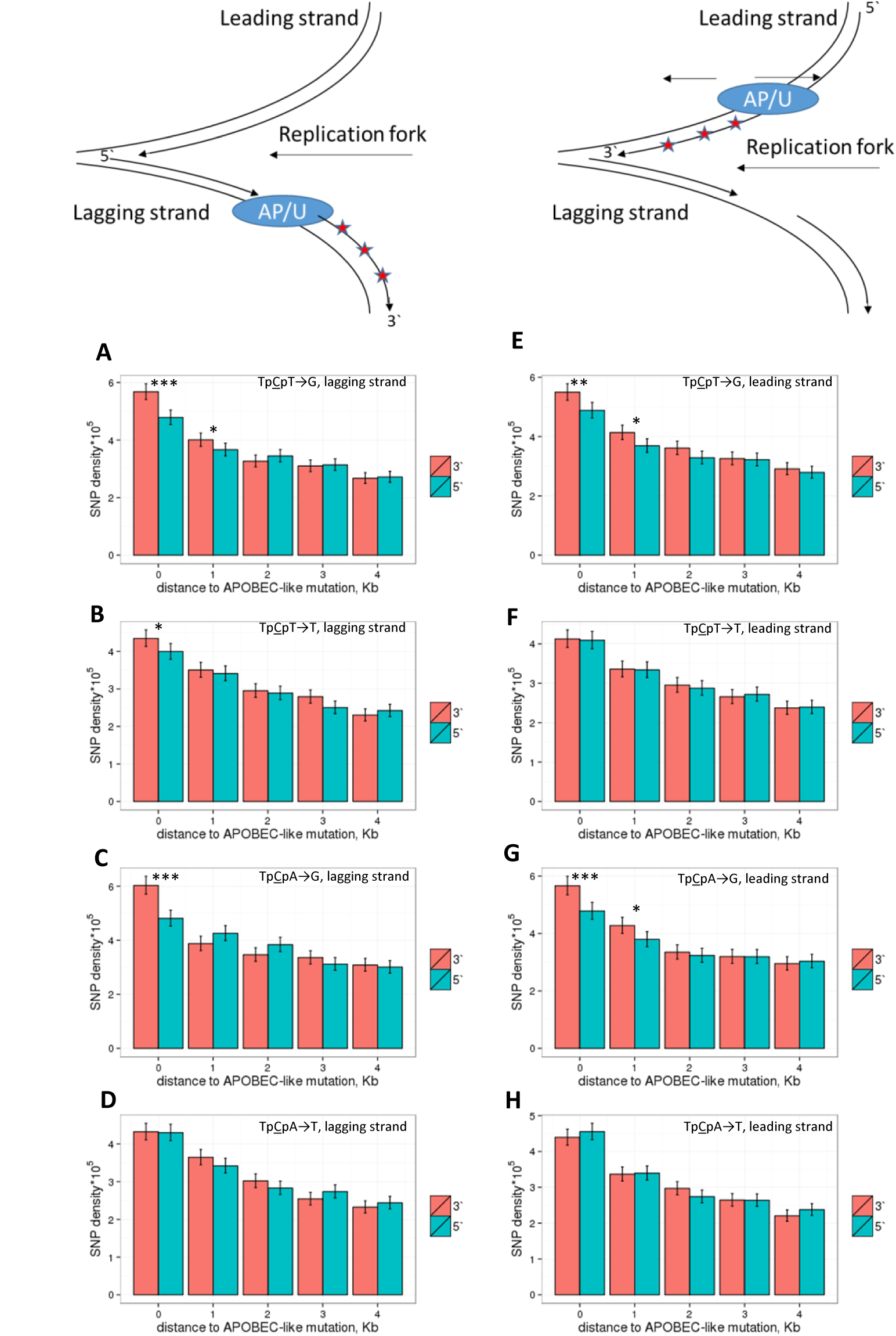
Density of linked SNPs 5’ and 3’ of a Tp<u>C</u>pW→K mutation. In the schematic depiction of a replication fork at the top of the figure, stars correspond to de novo mutations, and AP/U is the position of the Tp<u>C</u>pW→G mutation interpreted as the position of the abasic site, or the position of the Tp<u>C</u>pW→T mutation interpreted as the position of the uracil. The rate of mutations on the lagging (**A-D**) or the leading (**E-H**) strand is measured in 5 non-overlapping 1kb windows at increasing distance from the Tp<u>C</u>pW→K SNPs. ***, p < 0.001; **, p < 0.01; *, p < 0.05 (chi-square test).

A weaker effect was observed in one of the four comparisons for the C→T mutation (Figure 4B): a mutation rate asymmetry in the vicinity of Tp<u>C</u>pT→T mutations on the lagging strand (Figure 4B,D,F,H). This effect is particularly strong in the 400 nucleotides nearest to the C→T mutation (Figure S6). The Tp<u>C</u>pT→T mutations demonstrate the strongest bias towards the lagging strand, compared to the control mutation types (Figure 1), suggesting that APOBEC3A/B causes a high fraction of these mutations on the lagging strand. Therefore, our observations suggest that the mutation rate is increased 3’ of APOBEC3A/B-induced mutations. The difference between the mutation rates 5’ and 3’ of the mutation of interest is observed only for linked SNPs (Figure S7), implying that this association is due to the co-occurrence of mutations in a single mutational event rather than some underlying properties of the corresponding genomic regions.

The rate of all mutations is also strongly increased in the vicinity of a SNP in the same haplotype (Figure 4), in line with previous results^27^^−^^29^,^31^. This effect is weaker, and decays more gradually with distance between the mutations not linked with each other, compared with linked mutations (Figure S7).

## Discussion

Recent studies have linked most of the somatic mutations, mutations associated with cancers or with dedifferentiation of induced pluripotent stem cells with specific processes^10^,^11^,^32^. By contrast, only a minority of the heritable mutations are attributable to known mechanisms. Analyses of clustered mutations and L1 transposons have revealed a major role of low fidelity polymerase ζ and APOBEC3G in generation of heritable mutations^7^,^8^,^22^,^28^,^29^. Direct experiments have also uncovered the role of recombination in mutagenesis^33^,^34^. Still, the causes of many described and pervasive patterns observed in heritable mutations such as the cryptic variation of the site-specific mutation rate and heterogeneity of mutational spectra^35^^−^^37^ remain unknown.

Among mutation types, clusters of adjacent or nearby mutations can be attributed to specific mutational mechanisms most reliably, as chance occurrence of such clusters by conventional mechanisms is unlikely. However, the majority of mutations are dispersed, and few methods to study their origin are available. A type of mutation can be relatively easily linked to a specific mechanism only if its rate is unusually high, as is the case for <u>C</u>pG→T mutations.

Many mutations are associated with replication and preferentially occur at segments of DNA strand replicated as leading or lagging. A computational approach has been developed to discriminate between the leading and the lagging DNA strands^14^,^19^, providing a powerful tool to investigate strand-biased mechanisms that give rise even to dispersed mutations^14^,^19^,^38^,^39^. APOBEC3A/B-induced mutations, as well as mutations in mismatch repair (MMR)-deficient cells, are strongly biased towards the lagging strand^14^,^19^,^20^,^40^,^41^.

### Leading vs. lagging strand bias indicates that 20% of heritable mutations are induced by APOBEC3A/B

For all types of heritable single nucleotide substitutions, the asymmetry between the leading and the lagging strand is very weak^39^,^40^. Therefore, even a small admixture of mutations that are 2-3 times more prevalent on one of the strands can be detected (Figure 1), and the amount of such admixture can be estimated from the level of asymmetry. Previously^19^, found that *de novo* Tp<u>C</u>pW→K mutations obtained from^3^ are 1.38 times more frequent on the lagging strand than on the leading strand. In our analyses of the same dataset, the corresponding ratio is 1.23, but is not significantly different from 1 (p > 0.05). The difference between the results is likely due to differences in how fork polarity is measured. Still, the strand asymmetry of Tp<u>C</u>pW→K *de novo* mutations in a larger joined dataset^3^,^4^ yielded a significant 1.17-fold difference between strands (p-value=0.0174). This is in line with the ~1.15-fold asymmetry observed for rare SNPs. The observed ~15% asymmetry corresponds to a ~20% admixture of APOBEC3A/B-induced mutations. Still, the overall contribution of APOBEC3A/B to heritable mutagenesis is rather weak: we do not observe any tendency of C→K mutations to occur in the Tp<u>C</u>pW context among *de novo* mutations or in polymorphism data (Table S3), in line with^3^.

Our results show that knowledge of replication fork direction may be used to estimate the contribution of a specific process to heritable mutagenesis. We suggest that a similar set of approaches can be used to quantitatively estimate the fraction of APOBEC3G-induced mutations, especially among dispersed mutations, because recent experiments on bacteria showed that these mutations, too, are strand asymmetric^42^.

### Context dependence of heritable APOBEC3A/B-induced mutagenesis

The asymmetry between the lagging and the leading strand is stronger for mutations in the Tp<u>C</u>pT context than in the Tp<u>C</u>pA context independently of the mutation type (Figure 1). Notably, in all cohorts of cancers with a strong prevalence of APOBEC3A/B mutagenesis, C→G mutations also more frequently occur in the Tp<u>C</u>pT context than in the Tp<u>C</u>pA context (Table S4), in line with the accepted APOBEC3A/B signatures^10^.

Unexpectedly, the strand bias of the C→T mutation is also stronger in the Tp<u>C</u>pT context than in the Tp<u>C</u>pA context, even though APOBEC3A/B deaminates cytosines in the Tp<u>C</u>pA context slightly more frequently than in the Tp<u>C</u>pT context^16^,^17^. One reason for this discrepancy may be that the observed biases reflect repair rather than mutagenesis. The C→T mutation is a normal result of APOBEC-induced deamination giving rise to uracil followed by a U-A pairing during DNA replication^43^. Conversely, APOBEC-induced C→G mutations likely reflect initiated but unfinished repair and replication by low fidelity polymerases. This unfinished repair may introduce its own biases, which affect the context preferences of C→G mutations.

A higher fraction of APOBEC3A/B-induced mutations in the Tp<u>C</u>pT context in germline, reflected as a stronger asymmetry of both mutation types in the Tp<u>C</u>pT context, may arise due to the association between DNA repair and APOBEC activity. The activity of DNA repair systems provides substrate for APOBEC-induced mutagenesis^6^,^16^. Furthermore, somatic mutations in cancer that disrupt ERCC2 – a gene associated with nucleotide excision repair – are associated with a shift of APOBEC3A/B-induced mutagenesis to the Tp<u>Cp</u>A context^44^, supporting repair-dependent origin of mutations in the Tp<u>C</u>pT context. If the above reasoning is correct, this would imply that repair is associated with APOBEC3A/B-induced deamination in the Tp<u>C</u>pT context. If so, the stronger strand bias observed in the TpCpT context suggests that the majority of heritable APOBEC3A/B-induced mutations are associated with repair.

### Fraction of APOBEC3A/B-induced mutations is high in strand coordinated clusters

In cancer, APOBEC3A/B gives rise to mutational clusters spanning about 10Kb^12^,^14^. Linked SNPs, especially young ones, may be used to study such events, as mutational clusters observed in them have not yet been disrupted by recombination and are still detectible at distances of a few Kb^28^. Clustered mutations in the Tp<u>C</u>pW context are enriched in strand coordinated clusters by a factor of up to two. Moreover, for strand coordinated clusters, we observed a stronger bias towards the lagging strand than for dispersed mutations. Thus, the orthogonal approach based on strand coordination also detects the prevalence of APOBEC3A/B-induced mutations.

## Conclusion

APOBEC3A/B-induced mutations fuel cancer development, representing the second most prevalent mutational signature in it. Therefore, the cancer related activity of APOBEC3A/B should be slightly deleterious, although selection against it may be weak^45^. Here, we have shown that APOBEC3A/B also causes heritable mutations, thus increasing the mutation load. The fact that it is conserved by negative selection implies that its positive role in protection against retroelements or viruses outweighs its deleterious mutability. Thus, the maintenance of APOBEC represents a tradeoff between its advantageous function and deleterious mutagenesis.

## Material and Methods

### Mutational Data

The main set of results was obtained using polymorphism data from the 1000 Genomes Project^25^. We excluded exons and 10 nucleotides adjacent to each exon to reduce the contribution of selection, and only considered variants with low (< 1%) derived allele frequency, using the chimpanzee genome to determine the ancestral variant. Clusters were defined as pairs of SNPs of a particular type at distances between 10-5000 nucleotides from each other, such that at least half of the genotypes carrying the derived allele at one of the SNPs also carry the derived allele at the second SNP and vice versa. The same criteria were used to subdivide SNPs into linked (Figure 4) or unlinked (Figure S4) in analyses of SNP densities 5` or 3` of the considered SNP. For analyses of interspecies divergence, we used the human-chimpanzee-orangutan multiple alignment from the UCSC genome browser (https://genome.ucsc.edu/). We inferred substitutions in the human lineage after its divergence from the chimpanzee by maximum parsimony, using orangutan as the outgroup. Somatic mutations in cancers for whole genome sequences were obtained from^10^ and the TCGA consortium (https://tcga-data.nci.nih.gov/tcga/). Mutational clusters in cancers were determined as described in^14^, and mutational clusters of *de novo* heritable mutations obtained from trio data, as described in^3^. Cancers with a strong prevalence of APOBEC3A/B signature were defined on the basis of an increased rate of the Tp<u>C</u>pW→K mutation; this matches the definition of APOrich cancers in^14^.

### Leading vs. lagging strand asymmetry

The derivative of the replication timing at the position of the mutation was used as a proxy for the probability that the reference strand is replicated as leading or lagging in the current position, as described in^14^. The genome was categorized by these values into 6 (Fig. 2F-I) or 20 (Fig. 1 and Fig.2E) equal bins, with low value of the derivative corresponding to the propensity of the DNA segment to be replicated as lagging, and high value, as leading. For each bin, the numbers of substitutions and target sites were calculated. Each substitution was counted twice: as a substitution on the reference strand with the corresponding derivative of the replication timing, and as a complementary substitution with the inverse derivative. Thus, each plot of substitutions asymmetry (Fig. 1 and 2E-I) is symmetric with respect to zero. Confidence intervals were obtained for the relative risk in a 2x2 table. As a measure of the asymmetry, we used the ratio of the frequencies of complementary mutations between strands in the most extreme (first or last) bin, where the determination of the fp was the most confident. Tissue matched data on replication timing was used for analyses of asymmetry in cancers (MCF-7 for breast cancers, and IMR90 for lung cancers); this increased the observed level of asymmetry compared to our previous study and to a study by another group^14^,^19^. Since tissue matched replication timing for heritable mutations is unavailable, MCF-7 replication timing was used for this analysis.

### Estimation of the admixture of APOBEC3A/B mutations from the leading *vs* lagging strand asymmetry

Previously, using non-tissue specific replication timing data as in the current analysis of heritable mutations, we found that about two thirds of APOBEC3A/B mutations in cancer occurred on the lagging strand (Seplyarskiy et al. 2016). Therefore, the observed ratio of mutational frequencies on the lagging and on the leading strands (strand bias) *s* is
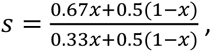
where *x* is the fraction of APOBEC3A/B-induced mutations among all mutations; 0.67*x* and 0.33*x* are the fractions of APOBEC3A/B-induced mutations on the lagging and on the leading strand respectively, with the coefficients 0.67 and 0.33 estimated from cancer data; and the coefficient 0.5 corresponds to the equal distribution of non-APOBEC3A/B-induced mutations between leading and lagging strands. Therefore,
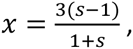
so that *s*=1.15 estimated for heritable mutations implies an admixture of APOBEC3A/B-induced mutations of 21%.

## Acknowledgements

We thank Wendy Wong, Shamil Sunyaev, Ruslan Soldatov and Nadezhda Terekhanova for useful discussion and Wendy Wong for help with data retrieving from^4^. This work was performed in IITP RAS and supported by the Russian Science Foundation (grant no. 14-50-00150)

## Contributions

V.B.S. performed analyses and convinced the study, all authors wrote manuscript, intensively discussed the study and contributed to design.

## References

1. Shendure, J. & Akey, J. M. The origins, determinants, and consequences of human mutations. Science 349, 1478–1483 (2015).

2. Kong, A. et al. Rate of de novo mutations and the importance of father’s age to disease risk. Nature 488, 471–475 (2012).

3. Francioli, L. C. et al. Genome-wide patterns and properties of de novo mutations in humans. Nat. Genet. 47, 822–826 (2015).

4. Wong, W. S. W. et al. New observations on maternal age effect on germline de novo mutations. Nat. Commun. 7, 10486 (2016).

5. Pfeifer, G. P. Mutagenesis at methylated CpG sequences. Curr. Top. Microbiol. Immunol. 301, 259–281 (2006).

6. Chen, J., Miller, B. F. & Furano, A. V. Repair of naturally occurring mismatches can induce mutations in flanking DNA. eLife 3, e02001 (2014).

7. Pinto, Y. et al. Clustered mutations in hominid genome evolution are consistent with APOBEC3G enzymatic activity. Genome Res. (2016). doi:10.1101/gr.199240.115

8. Knisbacher, B. A. & Levanon, E. Y. DNA Editing of LTR Retrotransposons Reveals the Impact of APOBECs on Vertebrate Genomes. Mol. Biol. Evol. 33, 554–567 (2016).

9. Montgomery, S. B. et al. The origin, evolution, and functional impact of short insertion-deletion variants identified in 179 human genomes. Genome Res. 23, 749–761 (2013).

10. Alexandrov, L. B. et al. Signatures of mutational processes in human cancer. Nature 500, 415–421 (2013).

11. Lawrence, M. S. et al. Mutational heterogeneity in cancer and the search for new cancer-associated genes. Nature 499, 214–218 (2013).

12. Roberts, S. A. et al. An APOBEC cytidine deaminase mutagenesis pattern is widespread in human cancers. Nat. Genet. 45, 970–976 (2013).

13. Burns, M. B., Temiz, N. A. & Harris, R. S. Evidence for APOBEC3B mutagenesis in multiple human cancers. Nat. Genet. 45, 977–983 (2013).

14. Seplyarskiy, V. B. et al. APOBEC-induced mutations in human cancers are strongly enriched on the lagging DNA strand during replication. Genome Res. 26, 174–182 (2016).

15. Burns, M. B. et al. APOBEC3B is an enzymatic source of mutation in breast cancer. Nature 494, 366–370 (2013).

16. Taylor, B. J. et al. DNA deaminases induce break-associated mutation showers with implication of APOBEC3B and 3A in breast cancer kataegis. eLife 2, e00534 (2013).

17. Chan, K. et al. An APOBEC3A hypermutation signature is distinguishable from the signature of background mutagenesis by APOBEC3B in human cancers. Nat. Genet. 47, 1067–1072 (2015).

18. Nik-Zainal, S. et al. Mutational processes molding the genomes of 21 breast cancers. Cell 149, 979–993 (2012).

19. Haradhvala, N. J. et al. Mutational Strand Asymmetries in Cancer Genomes Reveal Mechanisms of DNA Damage and Repair. Cell 164, 538–549 (2016).

20. Hoopes, J. I. et al. APOBEC3A and APOBEC3B Preferentially Deaminate the Lagging Strand Template during DNA Replication. Cell Rep. 14, 1273–1282 (2016).

21. Chan, K., Resnick, M. A. & Gordenin, D. A. The choice of nucleotide inserted opposite abasic sites formed within chromosomal DNA reveals the polymerase activities participating in translesion DNA synthesis. DNA Repair 12, 878–889 (2013).

22. Seplyarskiy, V. B., Bazykin, G. A. & Soldatov, R. A. Polymerase ζ Activity Is Linked to Replication Timing in Humans: Evidence from Mutational Signatures. Mol. Biol. Evol. 32, 3158–3172 (2015).

23. Rahbari, R. et al. Timing, rates and spectra of human germline mutation. Nat. Genet. 48, 126–133 (2016).

24. Terekhanova, Seplyarskiy, Soldatov & Bazykin. Evolution of local mutation rate and its determinants. Submited

25. 1000 Genomes Project Consortium et al. A global reference for human genetic variation. Nature 526, 68–74 (2015).

26. Aggarwala, V. & Voight, B. F. An expanded sequence context model broadly explains variability in polymorphism levels across the human genome. Nat. Genet. 48, 349–355 (2016).

27. Terekhanova, N. V., Bazykin, G. A., Neverov, A., Kondrashov, A. S. & Seplyarskiy, V. B. Prevalence of multinucleotide replacements in evolution of primates and Drosophila. Mol. Biol. Evol. 30, 1315–1325 (2013).

28. Harris, K. & Nielsen, R. Error-prone polymerase activity causes multinucleotide mutations in humans. Genome Res. 24, 1445–1454 (2014).

29. Zhu, W. et al. Concurrent nucleotide substitution mutations in the human genome are characterized by a significantly decreased transition/transversion ratio. Hum. Mutat. 36, 333–341 (2015).

30. Kochenova, O. V., Daee, D. L., Mertz, T. M. & Shcherbakova, P. V. DNA polymerase ζ-dependent lesion bypass in Saccharomyces cerevisiae is accompanied by error-prone copying of long stretches of adjacent DNA. PLoS Genet. 11, e1005110 (2015).

31. Schrider, D. R., Hourmozdi, J. N. & Hahn, M. W. Pervasive multinucleotide mutational events in eukaryotes. Curr. Biol. CB 21, 1051–1054 (2011).

32. Rouhani, F. J. et al. Mutational History of a Human Cell Lineage from Somatic to Induced Pluripotent Stem Cells. PLoS Genet. 12, e1005932 (2016).

33. Yang, S. et al. Parent-progeny sequencing indicates higher mutation rates in heterozygotes. Nature 523, 463–467 (2015).

34. Arbeithuber, B., Betancourt, A. J., Ebner, T. & Tiemann-Boege, I. Crossovers are associated with mutation and biased gene conversion at recombination hotspots. Proc. Natl. Acad. Sci. 112, 2109–2114 (2015).

35. Hodgkinson, A., Ladoukakis, E. & Eyre-Walker, A. Cryptic variation in the human mutation rate. PLoS Biol. 7, e1000027 (2009).

36. Johnson, P. L. F. & Hellmann, I. Mutation rate distribution inferred from coincident SNPs and coincident substitutions. Genome Biol. Evol. 3, 842–850 (2011).

37. Seplyarskiy, V. B., Kharchenko, P., Kondrashov, A. S. & Bazykin, G. A. Heterogeneity of the transition/transversion ratio in Drosophila and Hominidae genomes. Mol. Biol. Evol. 29, 1943–1955 (2012).

38. Baker, A. et al. Replication fork polarity gradients revealed by megabase-sized U-shaped replication timing domains in human cell lines. PLoS Comput. Biol. 8, e1002443 (2012).

39. Chen, C.-L. et al. Replication-associated mutational asymmetry in the human genome. Mol. Biol. Evol. 28, 2327–2337 (2011).

40. Andrianova, M., Bazykin, G. A., Nikolaev, S. & Seplyarskiy, V. Human mismatch repair system corrects errors produced during lagging strand replication more effectively. bioRxiv 45278 (2016). doi:10.1101/045278

41. Lujan, S. A. et al. Mismatch repair balances leading and lagging strand DNA replication fidelity. PLoS Genet. 8, e1003016 (2012).

42. Bhagwat, A. S. et al. Strand-biased cytosine deamination at the replication fork causes cytosine to thymine mutations in Escherichia coli. Proc. Natl. Acad. Sci. U. S. A. 113, 2176–2181 (2016).

43. Roberts, S. A. & Gordenin, D. A. Hypermutation in human cancer genomes: footprints and mechanisms. Nat. Rev. Cancer 14, 786–800 (2014).

44. Kim, J. et al. Somatic ERCC2 mutations are associated with a distinct genomic signature in urothelial tumors. Nat. Genet. (2016). doi:10.1038/ng.3557

45. Martincorena, I. & Campbell, P. J. Somatic mutation in cancer and normal cells. Science 349, 1483–1489 (2015).

